# Altered litter and cellulose decomposition across an anthropogenic habitat modification gradient in Sulawesi, Indonesia

**DOI:** 10.1101/2022.09.02.506341

**Authors:** Mukhlish J. M. Holle, Owen T. Lewis

## Abstract

Many tropical regions in Southeast Asia have experienced extensive habitat modification, creating a mosaic of forested and agricultural land. The capacity of these human-modified tropical landscapes to support biodiversity and ecosystem functions and services is of considerable practical interest.
Decomposition of plant material is a key process maintaining the nutrient cycle in both natural and agro-ecosystems, but few studies have documented the relative contributions of different guilds of decomposers, acting on different plant substrates, across different tropical land-uses.
We measured decomposition of leaf litter and cellulose across a gradient of anthropogenic habitat modification (forest, shrubland, and corn farmland) within Panua Nature Reserve, Sulawesi, Indonesia. The influence of fungi and the litter invertebrate community were investigated experimentally.
Decomposition of both substrates was significantly lower in corn plantations than in forest sites. Partial exclusion of litter invertebrates, but not fungi, significantly reduced decomposition, but the feeding guild composition of the litter invertebrate community did not differ significantly across habitat types.
Our results confirm that even small-scale agricultural areas adjacent to forest fragments can experience impaired ecosystem functions. These changes can be linked to reduced invertebrate activity, apparently independent of the functional structure of the litter invertebrate community. Effective management of human-modified landscapes will be needed to maintain nutrient cycling, even in areas where agroecosystems and tropical forests occur in close proximity.

**Highlights:**

- Habitat modification alters litter and cellulose decomposition
- Small-scale agriculture near forest fragments can impair ecosystem functions.
- Exclusion of litter invertebrates, but not fungi, significantly reduced decomposition.
- Maintaining nutrient cycling requires effective management of human-altered landscapes.

## Introduction

Tropical forests are a global priority for conservation and restoration because of their high biodiversity and provision of multiple ecosystem services (Brandon, 2014; Mokany, Westcott, Prasad, Ford, & Metcalfe, 2014; Brancalion et al., 2019; Strassburg et al., 2020). Southeast Asia is home to nearly 15% of the world’s tropical forests (Stibig, Achard, Carboni, Raši, & Miettinen, 2014; Estoque et al., 2019), but has the highest annual rate of deforestation and forest degradation among tropical regions (Mayaux et al., 2005; Miettinen, Shi, & Liew, 2011; Stibig et al., 2014). Reflecting global trends, conversion of forests to agriculture, especially oil palm and rubber, is the main cause of forest loss (Stibig et al., 2014) and the resulting biodiversity crisis (Wilcove, Giam, Edwards, Fisher, & Koh, 2013) in Southeast Asia. The fate of tropical forest biodiversity strongly relies on the effective management of human-modified landscapes, including areas where agroecosystems and tropical forests occur in close proximity (Gardner et al., 2009), and on the integration of management for ecosystem functions into land-use planning (Power, 2010).

Despite being a driver of tropical deforestation, agriculture production relies on ecosystem services that tropical forest ecosystems provide, such as soil fertility regulation through decomposition processes (Zhang, Ricketts, Kremen, Carney, & Swinton, 2007; Power, 2010; Gray & Lewis, 2014), which are mediated by rainforest biodiversity (Lewis, 2009). As agricultural expansion creates human-modified ‘matrix’ landscapes that now dominate most tropical regions (Brancalion, Melo, Tabarelli, & Rodrigues, 2013), protecting forest remnants may be crucial to support the fertility of nearby agricultural land.

Along with plant litter productivity (Veen et al., 2019; Giweta, 2020), decomposition is an important driver of soil nutrient availability through efficient cycling, especially where soils have low fertility (Yeong, Reynolds, & Hill, 2016; Grau et al., 2017). Furthermore, soil nutrients are also a limiting factor for tree seedling growth (Santiago et al., 2012). Decomposition of organic matter such as leaf litter and wood, is influenced by climatic conditions and biotic factors, such as the activities of decomposer communities: bacteria, fungi and macrofauna (including earthworms, termites and other litter invertebrates) (Power, 2010; Both, Elias, Kritzler, Ostle, & Johnson, 2017; Mohan, 2017). In general, decomposition is promoted by higher humidity and temperature (Salinas et al., 2011; Dan et al., 2016), a nutrient-rich environment (particularly for fungus-mediated decomposition) (Pascoal & Cassio, 2004), and a rich litter fauna, including both macro and microfauna (Gonzalez & Seastedt, 2001; Bradford, Tordoff, Eggers, Jones, & Newington, 2002). Numerous studies provide information on decomposition processes in tropical regions, but knowledge remains incomplete on two important aspects.

First, there is uncertainty about how decomposition rates vary across land-use gradients in interaction with biotic and abiotic factors. Studies have variously reported that decomposition rates in tropical forests are influenced by climate (Powers et al., 2009), habitat quality (e.g. litter: Kagezi et al., 2016; Yeong, Reynolds, & Hill, 2016, dung: Saleh, 2015), litter amount or quality (Ashford et al., 2013; Chellaiah & Yule, 2018), and the composition of the biotic community of arthropods (Ashford et al., 2013) or microbes (Chellaiah & Yule, 2018). However, there is a lack of consistency in the reported role of these different drivers. For example, Both et al. (2017) found that land use is a stronger driver than litter quality, while Powers et al. (2009) found that climate was the main driver, not mesofauna or litter type.

Second, there is a shortage of studies documenting and comparing decomposition of different substrates. While decomposition studies using leaf litter as the substrate are common, the study of wood decay has been relatively neglected (Pietsch et al., 2019). Cellulose-rich woody material is a key organic input, and is also important for tracking carbon balance changes including those linked to logging operations (Leitner, Davies, Parr, Eggleton, & Robertson, 2018). A recent study has shown that invertebrates such as insects have particularly pronounced effects on dead wood decomposition under the warm, humid conditions of tropical forests, accounting for a substantial fraction of the carbon flux from dead wood (Seibold et al., 2021).

In this paper, we investigate the importance of different components of the biota (invertebrates and fungi) as agents of decomposition across a gradient of land-use intensification in Gorantalo Province, Sulawesi, Indonesia. Our focal landscape is a mosaic of forest fragments, smallholder agriculture (dominated by corn, *Zea mays* L.) and abandoned agricultural areas where woody vegetation is regenerating. We explored decomposition of different types of plant matter (leaf litter and cellulose), compared decomposition rates across habitat types, and manipulated the composition of the decomposer community (with fungi excluded or included). We also measured the abundance and community composition of the litter invertebrate community associated with our measures of decomposition. We predicted that rates of decomposition would be higher in less-modified habitats, and that the biotic and abiotic factors influencing decomposition would differ among substrates, with both fungi and litter invertebrates playing a role.

## Materials and Methods

### Site description

The research was conducted from 15^th^ February – 27^th^ April 2019 in Panua Nature Reserve (PNR), which is situated in the Pohuwato Regency (0° 31 59 N, 121° 49 30 E), Gorontalo Province, Sulawesi, Indonesia, and protected under Indonesian environmental legislation because of its biotic uniqueness. However, human disturbance, especially forest clearance for agriculture, is significant and ongoing within the reserve (Nursafingi & Holle, in review). Since 1980, forest cover in Panua Nature Reserve has been reduced by 6582 hectares, equivalent to 13.4% of the total reserve area, with an estimated annual deforestation rate of 162 hectares per year. The main deforestation driver is conversion to agriculture, which reduced the forested area by 5704 hectares (Nursafingi & Holle, in review). Panua Nature Reserve has an annual rainfall of 1500-2000 mm, with five or six consecutive wet months and three or fewer consecutive dry months (Whitten, Mustafa, & Henderson, 1987); the period of fieldwork corresponded to the rainy season. The study was conducted in the major habitats within the nature reserve: forests, shrublands and corn fields. We focused on human-modified forest areas within 100 m of the forest edge, subjected to varying levels of ongoing small-scale tree-cutting and adjacent to farmland. Forests of this type, rather than less-disturbed areas deeper into the protected area, are typical of the agriculture/forest matrix in human-modified landscapes of SE Asia. Forest sites had experienced low-intensity selective logging of large and emergent trees, but retained closed-canopy structure. Shrubland sites were dominated by an invasive plant species, *Chromolaena odorata* (L.) R.M. King & H. Rob. growing to 2 m tall, which produces allelochemical compounds that inhibit other plants (Laxman, Desai, & Krishna, 2019). These shrublands are formed due to abandonment after forest clearing, or when cropped corn fields are left fallow after the planting season. Cornfield sites are managed by smallholder farmers who have illegally cleared land for agriculture within the nature reserve. Sampling in the cornfields sites was started three weeks after planting, when corn stalks started to create locally-shaded microclimates, allowing the study to be completed before corn was harvested approximately three months after planting. Shrubland and corn farming are both non-permanent land uses: outside of the planting season (mostly the wet season), fallow corn farmland can develop into either shrubland, bare ground, or grassland, while some shrublands are periodically cleared to plant corn.

### Study designz

Decomposition experiments used two different substrates, leaf litter and cellulose, representing natural sources of organic matter on the forest floor. We measured litter decomposition, cellulose decomposition and the litter invertebrate community concurrently in different habitat types at two-week intervals.

We established a study design of seven spatial blocks, with each habitat type (forest, shrubland or corn farmland) represented once within each block. Individual blocks were widely separated from each other (by distances of 0.7-18 km), ensuring appropriate replication (Figure 1). In six of the blocks, sites representing the three habitat types occurred within close proximity (less than 0.5 km apart), while in the seventh block the corn farmland site was located 18 km from other habitats for practical reasons. Each habitat replicate comprised a 20×20 m plot.

**Figure 1.**
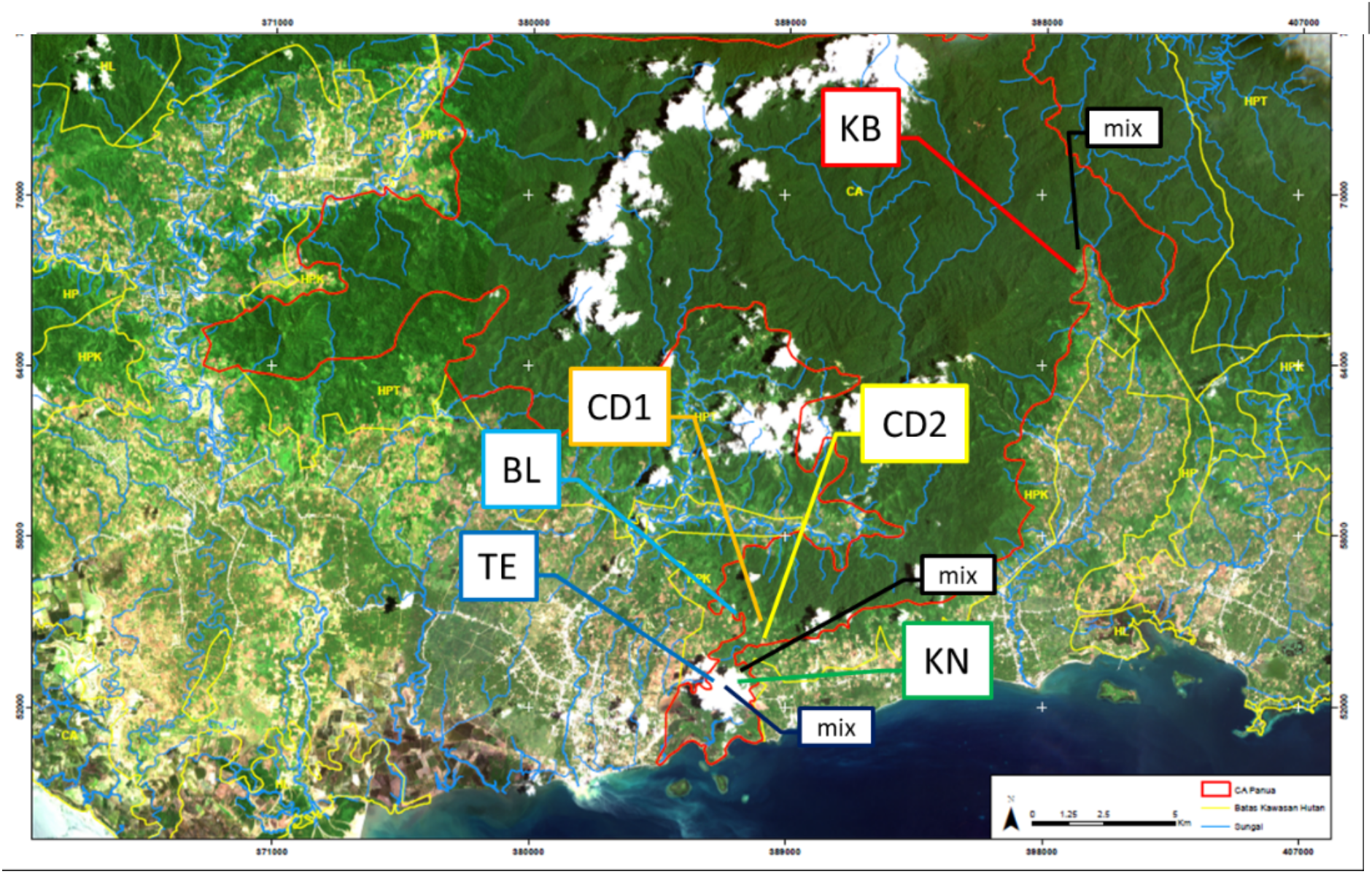
Map showing the blocked distribution of study sites within the human-modified tropical landscape of Panua Nature Reserve (reserve boundary shown in red). Block labels in the different box line colour refer to location names (KB: Karya Baru; BL: Bulangita; CD1: Cek Dam 1; CD2: Cek Dam 2; TE: Teratai; KN: Kebun Nurdin). ‘Mix’ is a mixed block where sites in different habitats are separated more widely (Image source: Burung Indonesia Gorontalo).

**Figure 2.**
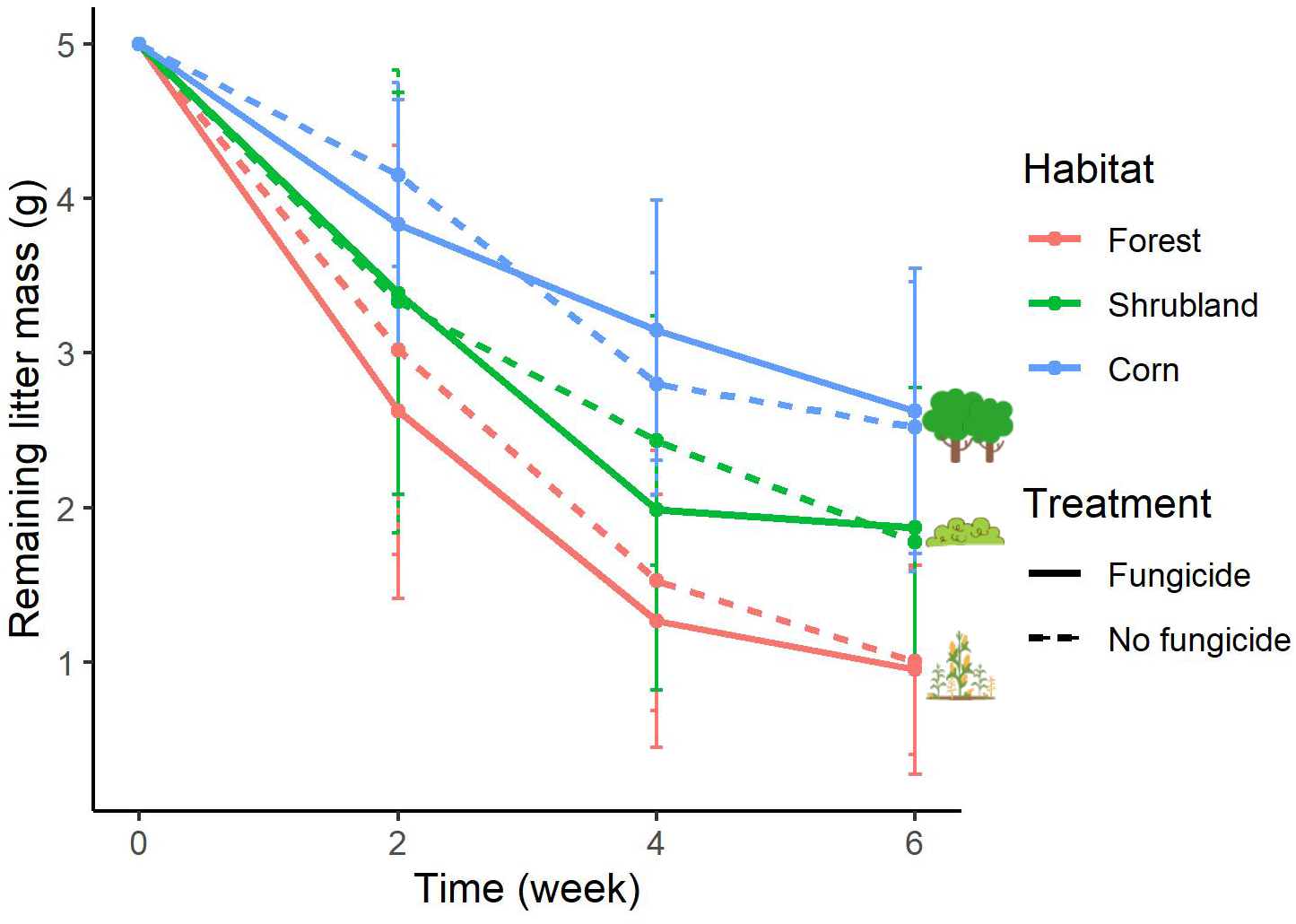
Remaining mass of *Macaranga* sp. leaf litter in three habitats measured every two weeks (n=689). Error bars show standard errors of the mean.

### Leaf litter decomposition and litter invertebrate extraction

In each block, 108 litter bags were deployed, making a total of 756 litter bags: 7 blocks x 3 habitats (forest, shrubland, corn) x 2 treatments (fungicide, control) x 3 exposure times (2, 4, 6 weeks) x 6 replicates. Litter decomposition rates were calculated adapting methods from Powers et al. (2009). Litter was derived from fresh whole-leaves of *Macaranga* sp. trees following previous studies (Plowman & Ewers, 2012; Chellaiah & Yule, 2018); these pioneer trees are frequent but not dominant at all our study sites, making it unlikely that there will be a home-field advantage, and their leaves decompose quickly (within a month: Chellaiah & Yule, 2018). Although naturally-abscised, senescent leaves might more accurately reflect the chemistry of natural substrates, we used dried fresh leaves for our experiment, consistent with previous decomposition experiments (Moore & Fairweather, 2006; McLaren & Turkington, 2010) and because it was not feasible to collect sufficient freshly-abscised leaves. Collected leaves were exposed in full sun for three days until completely dry, indicated by their crisp texture. Larger dried leaves were broken up by hand into approximately 6 x 6 cm fragments that could be inserted into the litter bags. Each 10 x 15 cm non-degradable litter bag was filled with a 5-g sample of litter and sealed using rust-proof clips. Litter bags with coarse mesh (0.5 cm in diameter) were used to allow access to micro-, meso- and macro-fauna (Bradford et al., 2002).

In the fungus suppression treatment, litter bags containing litter were dipped in liquid fungicide (0.25 ml/L of SCORPIO 250 EC Fungicide, active ingredient difenoconazole) immediately before being placed on the ground. Controls were untreated. Sets of six litter bags were tied to a 1-meter length of cord to facilitate relocation. The distance between sets was 13 meters (Powers et al., 2009). Thirty-six litter bags per block (252 litter bags in total) were collected (two litter bags in every set every two weeks) for extraction of litter-dwelling invertebrates to assess decomposition rate changes through time in parallel with litter mass measurement. Particular care was taken when retrieving bags to minimise loss of smaller fragments through the mesh. Inevitably, some litter will be lost from coarse-meshed bags without being decomposed, but any such effect will have been consistent across treatments.

Immediately after litter bag retrieval from the field, litter was carefully removed from the bag and immediately placed in an open-topped plastic bag to ensure that no invertebrates escaped, and to avoid overheating. Samples were then transferred to Tullgren funnels as soon as possible to extract invertebrates while drying the leaf litter. The contents of each collected litter bag were exposed separately under a 15 Watt light bulb for approximately 12 hours. Extracted invertebrates were preserved in 70% ethanol. Under a digital microscope, each individual invertebrate with body size > 2 mm was counted, identified to higher taxon level, and categorised in terms of its ecological role (decomposer, omnivore, and predator) on the basis of feeding guilds. Within feeding guilds, invertebrate samples were assigned to taxon groups, mostly to Order level, but in some cases to Family (Formicidae) or Class (Chilopoda, Diplopoda). This approach of assigning taxon groups to feeding guilds is likely to be imperfect because litter fauna from the same taxon do not always occupy the same feeding guild (Ashford et al., 2013), but further identification and documentation of feeding guilds was not practicable in this species-rich and poorly-studied community within the scope of the current project. Instead, taxa which can include species within multiple feeding guilds, such as ants, were categorized into the generalist feeding guild of omnivores. Dry litter mass was measured using a digital balance after soil particles, clinging roots, and other debris were removed by hand sorting.

### Cellulose decomposition experiment

As a standardised material that represents wood, cellulose baits (single ply, unscented coreless toilet rolls) were deployed to measure cellulose degradation in various habitats for two months from 15^th^ February – 27^th^ April 2019. Toilet rolls have been used previously to assess termite-mediated decomposition (e.g. Leitner et al., 2018). These cellulose baits were located at the same sites as the litter bag experiment. In total 126 toilet rolls (7 blocks x 3 habitats x 6 replicates per block) were used. The toilet rolls were not dried initially. They had very low water content, were used directly after removal from plastic packaging, and varied little in initial mass. Recollected toilet rolls were dried in direct sun for three days until completely dry, indicated by their constant dry mass. Cellulose decomposition was calculated by subtracting remaining toilet roll mass from the initial toilet roll mass (33.89 g ± 0.20 (SD)).

### Data analysis

Decomposition data were expressed as decomposed litter mass and as a decomposition rate (k), following Guendehou et al. (2014), to evaluate the decomposition process and to measure the speed at which leaf litter is broken down by decomposers respectively. Decomposition rate (k) was estimated using a negative exponential equation:

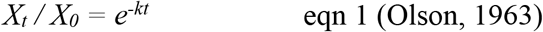

where *X_t_* is litter mass at time *t*, and *X_0_* is original litter mass, and *X_t_ / X_0_* is therefore the fraction of the original litter mass remaining at time *t*. Where litter was completely decomposed, litter mass values of zero were input as 0.01g to allow *k* value calculations. After calculating k values for the seven replicate sites of each habitat, we used linear mixed-effects models to assess the effects of habitat type on decay rates, with block as a random effect. The *k* values were log-transformed for these analyses as heteroscedasticity was detected in the standardised residuals.

All statistical analyses and graph plotting were performed using R (R Core Team, 2021). Analyses used data from all blocks with the exception of analyses of litter invertebrates, for which data were missing for the KB block. The effects of time of incubation, habitat type and invertebrate community on decomposed litter mass were analysed using linear mixed effects models (using the *lme* function in the *lme4* or *nlme* package) with habitat type, fungicide treatment and litter invertebrate abundance as fixed factors and block as a random effect. Full models with interactions were fitted, and then non-significant interaction terms dropped from the model until the minimal adequate model was obtained according to the AIC scores. The minimal adequate model was the best model as judged by AIC, so the effects of explanatory variables on decomposed litter mass were reported using this model. Analysis of variance (ANOVA) was conducted to compare means among groups. To identify the groups that differed significantly, linear mixed-effects model analysis was followed by post-hoc analysis with Tukey Contrasts test using the *glht* function in the *multcomp* package in R.

To compare decomposition of leaf litter and cellulose we analysed the effects of habitat type on decomposition using data from week 6. Linear mixed effects models (using the *lme* function in the *lme4* or *nlme* package) were used to test the effects of habitat type and fungicide as fixed factors and block as a random effect (Bates et al., 2015). Analysis of Similarity (ANOSIM) (*anosim* function in the *vegan* package) was used to test for statistical differences among the invertebrate communities of the three habitat categories and two fungicide treatment categories (Dixon, 2003; Oksanen, 2022). Analyses focused on invertebrate relative abundance data obtained from the litterbags extracted at week 2 of the experiment, to avoid underestimating abundances; rapid decomposition meant that leaf litter mass was much reduced in later weeks, likely limiting invertebrate colonisation and occupancy. Sampling points where no invertebrates were recorded were omitted from the analysis. Data were visualised using a Non-metric Multi-dimensional Scaling (NMDS) plot, generated in *ggplot* using the Bray-Curtis distance measure.

## Results

### Effects of habitat, fungicide, and time on litter decomposition

Remaining litter mass decreased significantly with time (F_1, 698_=364.03, p <0.001), and differed significantly among habitats (F_2, 698_=130.76, p <0.001). Remaining litter mass was lowest in forest, highest in corn, and intermediate in shrubland, with all pairwise comparisons statistically significant (Tukey Contrasts test). There was no significant effect of fungicide on litter decomposition (F_1, 698_=6.71, p=0.43) (Table 3). Although the interaction between fungicide treatment and time was significant (F_1, 671_=4.49, p <0.03, linear mixed effects model), the best-fitting model on the basis of comparing AIC values did not include the interaction term.

Similarly, decomposition rates measured using *k* values differed significantly across habitat types (F_2,11_=12.823, p=0.0013). Rates were highest in forest (0.293±0.03 (SE)), lowest in corn (0.127±0.01 (SE)), and intermediate in shrubland (0.224±0.02 (SE)), with all pairwise comparisons statistically significant (Tukey Contrasts test) except the comparison between forest and shrubland (Figure 3).

**Figure 3.**
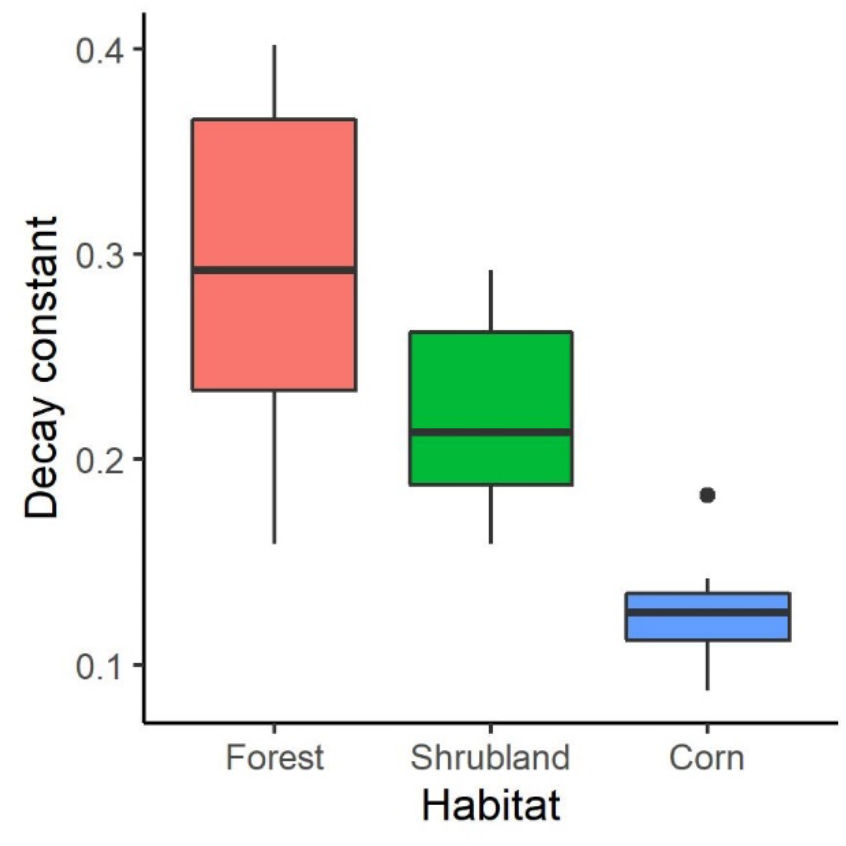
Decay constant (k) of *Macaranga* sp. leaf litter in three habitats (n=20 sites spread across 7 blocks; data missing for one Shrubland site). The boxplots show the median (central bar), upper and lower quartiles, and whiskers including points within 1.5 interquartile range. Dots outside of the boxplots are outliers.

### Comparison of decomposition of leaf litter and cellulose across habitat types

Analysing litter decomposition in the final retrieval at week 6 (Figure 4), the mass of litter remaining differed significantly among all habitats (Tukey contrasts test). Decomposition was highest in forest, lowest in corn plantations, with shrubland intermediate (F_2, 212_=78.62, p <0.001). There was no significant effect of fungicide (F_1,212_=0.71, p =0.39). Cellulose degradation differed significantly among habitats (F_2,89_=25.54, p <0.001), with mass remaining significantly lower in forest than shrublands and corn, which did not differ significantly from each other (Tukey Contrasts test).

**Figure 4.**
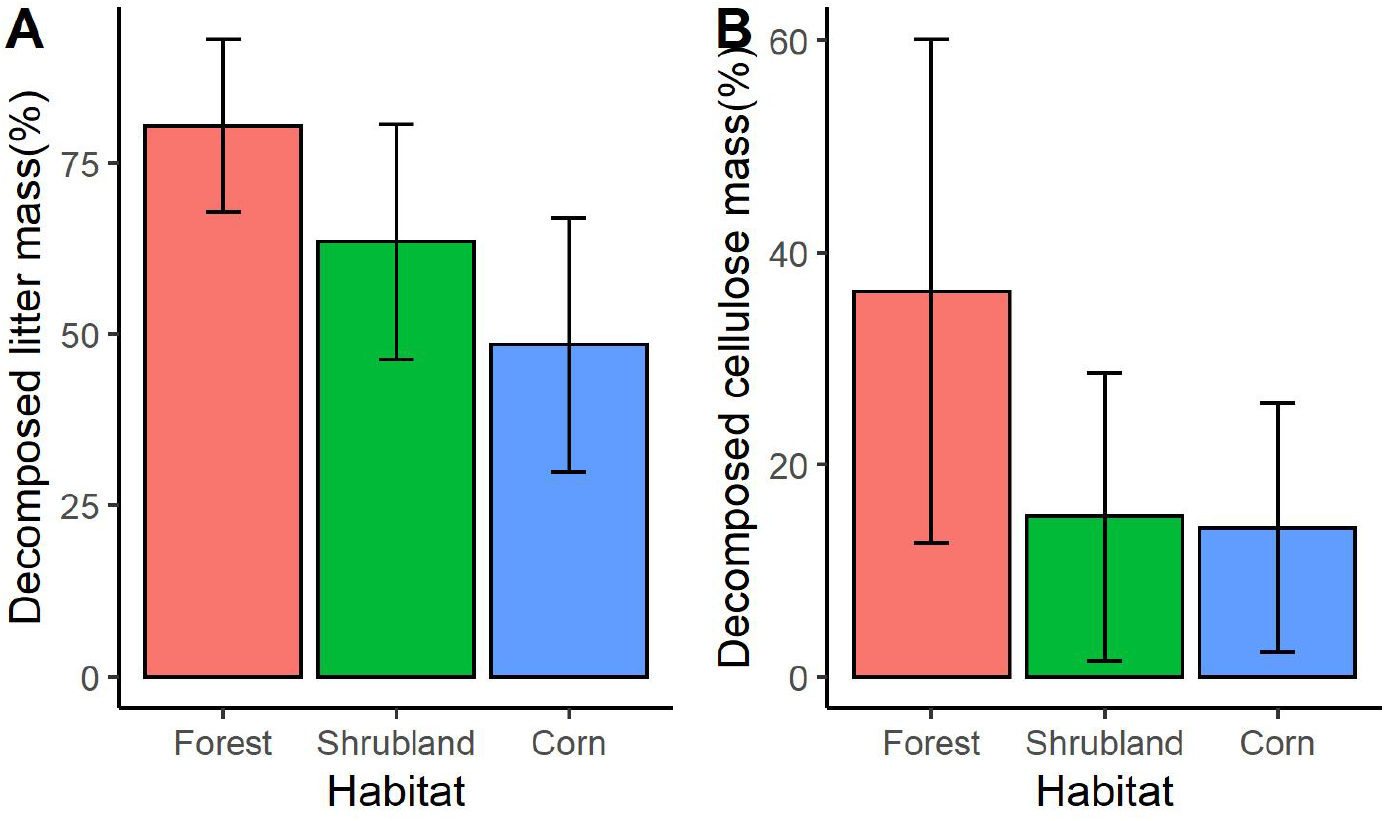
Decomposed mass of (A) leaf litter (n=222) and (B) cellulose (n=98) exposed in three habitat types and retrieved after six weeks. Visualisation in (A) disregards the fungicide treatment which had no significant effect on decomposition. Error bars show standard errors of the mean.

**Figure 5.**
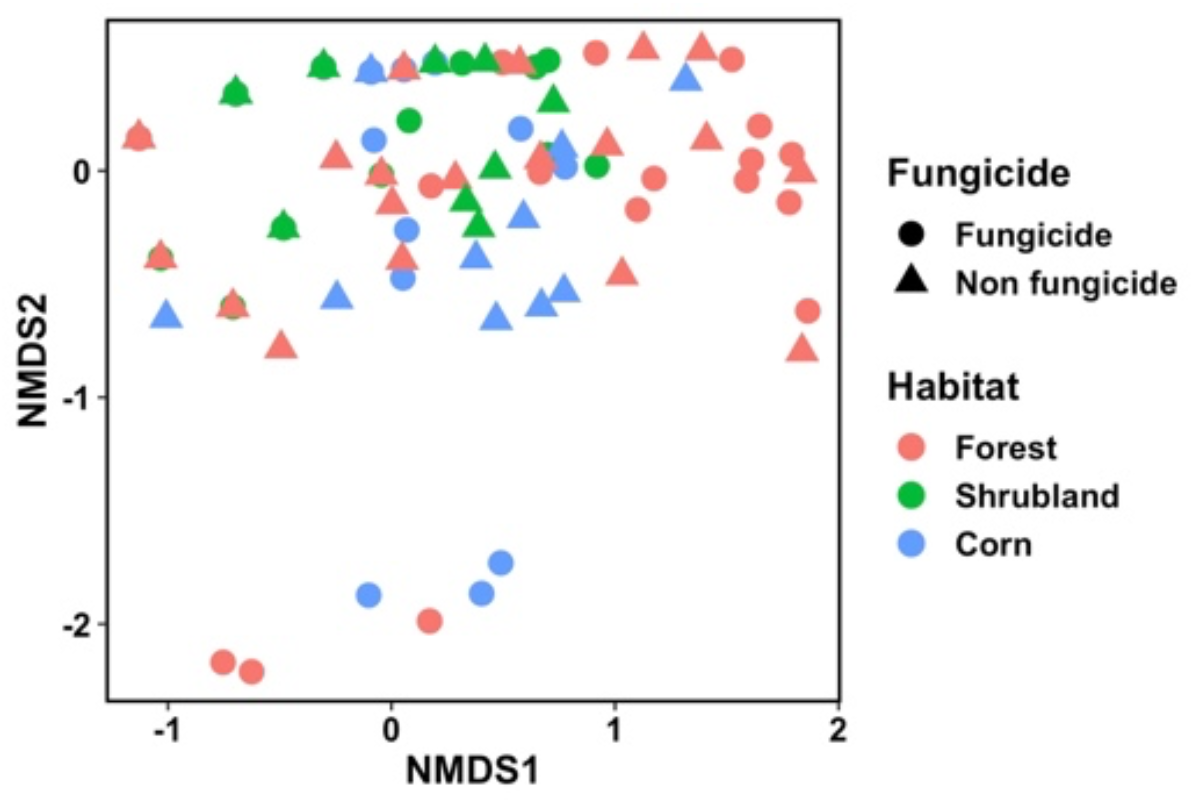
Non-metric multidimensional scaling (NMDS) of litter invertebrate community composition in relation to habitat type and fungicide treatment for 2 weeks litterbag exposure. Each habitat type is represented by a different colour and the fungicide treatment is indicated by symbol shape. The Bray-Curtis dissimilarity index was used for determination of dissimilarities among litter invertebrate communities. Stress value is 0.1148, indicating good fit of the model.

### Litter invertebrate abundance in different habitats and through time for different feeding guilds

We identified 3572 individual litter invertebrates in total. Total abundance was highest in forest sites (1571 individuals) followed by corn farming (1093 individuals) and shrubland (906 individuals). Total abundances of decomposers, omnivores, and predators were 694 (19.42%), 2763 (77.35%), and 115 (3.22%) individuals, respectively, which differed significantly. Based on the week 2 relative abundance data, there was a statistically significant difference in litter invertebrate community abundance between habitats (ANOSIM, R=0.028; p=0.0356). No significant fungicide effects on invertebrate abundance were detected in any week (ANOSIM, p>0.1).

## Discussion

Our study documents the effects of habitat modification on multiple dimensions of decomposition in a tropical setting, alongside changes to the litter invertebrate community. We found that habitat modification altered decomposition consistently for both leaf litter and cellulose substrates, with reduced rates in more modified environments.

### Effects of habitat modification on decomposition of plant substrates

Litter decomposition was most rapid in the forest sites, lower in shrubland, and lowest in corn farmland. This is consistent with previous studies that have found lower decomposition rates in agricultural sites compared with natural or less-modified sites in tropical regions (Didham, 1998; Attignon et al., 2004; Kagezi et al., 2016), especially at low elevations (Becker & Kuzyakov, 2018). Land-use change from tropical forest to intensively managed agricultural land has been shown to affect functional trait composition; this could impair the functional stability of litter invertebrate communities, with managed monoculture plantation tending to have more inconsistently assembled and compositionally unstable communities (Mumme et al., 2015).

Higher decomposition rates are to be expected in closed canopy-sites, where litterfall input will be high (Paudel, Dossa, Xu, & Harrison, 2015), supporting high decomposer biomass and activity. Microclimatic conditions in these habitats are also expected to favour decomposer activity (Lodge, Cantrell, & González, 2014). Like leaf litter decomposition, cellulose decomposition was higher in the forest habitat. Our corn farming and shrubland sites experience negligible natural inputs of woody material, so low prevalence of cellulose-adapted taxa in these habitats is likely to have limited the capacity of decomposer communities to process cellulose baits. Wood and cellulose decomposition in tropical rainforests is often mediated by termite activity (Jouquet, Traoré, Choosai, Hartmann, & Bignell, 2011; Leitner et al., 2018; Ashton et al., 2019). However, termites were not the sole decomposers of cellulose baits in our experiment, with large-bodied millipedes, specifically *Salpidobolus* sp., also having an important role (JH, personal observation). The main correlates and drivers of wood decomposition rates remain uncertain. For example, while some studies show that wood decomposition is more influenced by temperature than by tree species and decomposer diversity (e.g. Pietsch et al., 2019) others have concluded differently (e.g. Dossa, Paudel, Cao, Schaefer, & Harrison, 2016).

### The absence of fungicide effects on leaf litter decomposition

Surprisingly, fungicide treatment did not slow leaf litter decomposition or lower decomposition rates in any of the three habitats. At face value, this suggests that the contribution of fungi to decomposition in our system is small compared to that by litter invertebrates. There are four alternative explanations for this result. First, our methods may have been ineffective in suppressing fungal activity, perhaps because we used low fungicide concentrations corresponding to the minimum recommendations of the manufacturer, or because the active ingredient in our fungicide, difenoconazole, had low effectiveness (Masiello, Somma, Ghionna, Francesco Logrieco, & Moretti, 2019). A previous study found that the azole fungicide tebuconazole had no significant effect on leaf consumption by the aquatic amphipod crustacean *Gammarus fossarum* unless at very high concentrations (500 μg/L) (Elskus et al., 2016). Second, fungus-mediated decomposition is improved by nutrient enrichment (Pascoal & Cassio, 2004), and fungal decomposition activity is decreased as a result of the low-nutrient content in tropical soils (Yeong et al., 2016). A third possible mechanism is that, as the study took place during the rainy season, the fungicide treatment may have been ineffective because of rapid leaching during rainfall events (Lorenz et al., 2017). Finally, the role of fungi in decomposing leaf litter could be taken over by other taxa, such as invertebrates. Macrofaunal contributions to leaf litter decomposition can range widely, from 1.6% to 66% depending on litter type and climatic conditions (Slade & Riutta, 2012). The *Macaranga* sp. leaves we used do not contain high amounts of heavier structural compounds such as lignin, cellulose and fibre which are associated with high leaf tensile strength (Chellaiah & Yule, 2018). Thus, *Macaranga* leaves may be relatively susceptible to invertebrate feeding and mechanical fragmentation.

We grouped litter invertebrates into three feeding guilds: decomposers, omnivores, and predators. Categorizing invertebrates by their functional group can be a robust approach for assessing ecosystem conditions (Cummins, Merritt, & Andrade, 2005; Menezes, Baird, & Soares, 2010) and evaluating impacts of land use change (Schulze et al., 2004). Invertebrate abundance responded idiosyncratically among different habitat types. Litter invertebrate relative abundance was different among habitat types and was reduced in human-modified corn and shrubland. This is consistent with previous studies documenting that land use change from forests to agriculture altered invertebrate communities in both tropical (Barnes et al., 2014; Edwards et al., 2012; Jones et al., 2003) and temperate ecosystems (Eggleton et al., 2005). Resource availability might drive this pattern, with forest habitat providing greater and higher-quality food for litter invertebrates (Richardson et al., 2005). The more leaf litter, the more litter invertebrates are available to colonise litter bags. Alongside reduced litter abundance, reduced litter diversity could also contribute to these patterns. Litter diversity is correlated with vegetation diversity, which inevitably declines when diverse tropical forests are modified or converted to agricultural monocultures (Gillison et al., 2003). Nonetheless, existing data suggest that litter quantity has a greater effect on litter invertebrates than litter diversity (Lu et al., 2016).

Consistent with the result for decomposition, fungicide treatment did not affect the litter invertebrate community significantly in our experiment. While some studies have demonstrated that fungicides negatively affect invertebrate communities (Fernández et al., 2015; Flores et al., 2014), others report contrasting results, suggesting that fungicide toxicity is variable or context dependent. Fungicide concentrations may not have exceeded the tolerance range of litter invertebrates in our study, and higher concentrations might be expected to have more pronounced impacts, as documented in previous studies (French & Buckley, 2008; Zubrod et al., 2017). Furthermore, long-term and repeated exposure to low fungicide concentrations could increase chronic effects (Elskus et al., 2016).

## Conclusion

Habitat modification markedly reduced decomposition rates of both litter and cellulose. Across all habitat types our data are consistent with a minor role for fungi in decomposition of these substrates, relative to litter fauna. Studies quantifying the contribution of litter decomposition to nutrient inputs in the soil, and further investigating how decomposer biodiversity affects decomposition will be needed to better understand the effects on land-use change on ecosystem services mediated by decomposers, as well as the biotic and abiotic mechanisms involved.

## Acknowledgements

We are grateful to BKSDA Gorontalo, Ministry of Environment and Forestry of Indonesia for research permissions. Our work was approved by the Ministry of Environment and Forestry of the Republic of Indonesia, under permit 46/BKSDA.Sulut/TU/SIMAKSI/XI/2018. Staff of BKSDA Gorontalo (Tatang Abdullah, Abdul Sadam, and Rosalia Nurma) assisted field data collection. Staff of Burung Indonesia (Pantiati and Afi Nursafingi) supplied site information and helpful discussion. Annisa Firdaus assisted sample preparation and identification. Chris Terry, Fujinuma Junichi, and Sumali Bajaj helped with data analysis. MJMH is financially supported by LPDP Scholarship from Ministry of Finance of Indonesia. We thank all CERO members for their constructive comments.

